# The synchronization and adaptation of *Neurospora crassa* circadian and conidiation rhythms to short light-dark cycles

**DOI:** 10.1101/351189

**Authors:** Huan Ma, Luyao Li, Jie Yan, Yin Zhang, Weirui Shi, Yunzhen Li, Menghan Gao, Siyu Pan, Ling Yang, Jinhu Guo

## Abstract

Circadian clocks control the physiological and behavioral daily rhythms to adapt to the changing environment with a period of ~24 h. However, the influence and mechanism of extreme light-dark cycles on the circadian clock remain unclear. We show that, in the fungus *Neurospora crassa* under short LD cycles, both the growth rate and the ratio of microconidia production contributes to adaptation in LD12:12 (light for 12 h and dark for 12 h, periodically). Mathematical modeling and experiments demonstrate that in short LD cycles, the expression of the core clock protein FREQUENCY is entrained to the LD cycles when LD>3:3 while it free runs when *T*≤ LD3:3. We investigated the changes in circadian/diurnal rhythms under a series of different LD conditions, and the results showed that conidial rhythmicity can be adapted to the short LD cycles. We further demonstrate that the existence of unknown blue light photoreceptor(s) and the circadian clock might promote the conidiation rhythms that resonate with the environment. A high-intensity light induced the expression of a set of downstream genes involved in various metabolic pathways. The ubiquitin E3 ligase FWD-1 and the previously described CRY-dependent oscillator system were implicated in regulating conidiation under short LD conditions.

---

The Earth rotates with a period of approximately 24 h, which results in the periodic change in of many environmental factors. Circadian clocks are the inner mechanisms that allow organisms to adjust their physiology and behavior to the daily cycling of environmental factors, e.g., light and temperature (Bell-Pederen *et al*. 2005).

In certain ranges, non-24h light/dark cycles can entrain the circadian rhythms, and the flexibility of this entrainability dramatically varies among different organisms. Within the range of entrainment which is also called the limit of stable entrainment, the rhythms change in accordance with the non-24 T-cycles and show stable phase angles; beyond the ranges, the circadian clocks free-run (Refinetti 2004; Madrid *et al*. 1998; Boulos *et al*. 2002; Abraham *et al*. 2010; Jewett *et al*. 1994). The entrainment ranges have been characterized in a number of species including human, hamster and Neurospora (Ralph and Menaker 1988; Nsa *et al*. 2015; Dong *et al*. 2008; Dunlap *et al*. 2004; Binkley 1990; Czeisler *et al*. 1999). In the wild-type filamentous fungus *Neurospora crassa*, the conidiation banding rhythms are consistent with a range of T-cycles that include LD6:6, LD12:12 and LD14:14, which illustrates the masking effects (Nsa *et al*. 2015). These observations have been extended to conidiation rhythms in LD3:3 and LD9:9 (Dong *et al*. 2008).However, under extremely short T-cycles that exceed the thresholds, the rhythms free run in certain species, including Neurospora, *Canavalia ensiformis* and sparrow (Dunlap *et al*. 2004; Binkley 1990).

It is very challenging for the endogenous circadian rhythms to be entrained to non-24 h LD cycles compared to the rest-activity rhythms (Czeisler *et al*. 1999; Lewis and Lobban 1957). Under such conditions, the endogenous rhythmicities may desynchronize with the behavioral rhythmicities, which leads to inadaptation or disorders that have been observed in a number of checked organisms (Czeisler *et al*. 1999; Ouyang *et al*. 1998; Highkin and Hanson 1954; Hut and Beersma 2011).

Neurospora is an important model organism for circadian clock study; it exhibits overt clock phenotype in its conidiation. On the molecular level, the core circadian clock system of Neurospora is composed of two positive elements: White Collar 1 (WC-1) and WC-2 and one negative element: FREQUENCY (FRQ). The FRQ/WC-based circadian oscillator (FWO) dictates the circadian rhythms at molecular and physiological levels (Hurley *et al*. 2015; Baker *et al*. 2012). WC-1 harbors three PER-ARNT-SIM (PAS) domains; it is responsible for light responses of circadian clock as a blue light photoreceptor (Froehlich *et al*. 2002; He *et al*. 2002).

Neurospora has several additional validated or putative photosensing-associated factors, including VVD, CRY, NOP-1, PHY-1 and PHY-2 (Schwerdtfeger and Linden 2001; Corrochano 2007). VIVID (VVD), another PAS domain-containing protein, is a flavin-binding blue-light photoreceptor that regulates the light responses and temperature compensation of the Neurospora circadian clock (Zoltowski *et al*. 2007; Heintzen *et al*. 2001). Interestingly, *acry-dependent oscillator gate-1* (*cog-1*) mutation led to conidiation rhythmicity in constant light in Neurospora, suggesting a role for *cog-1* in regulating light sensing in the circadian clock. The *cog-1* related oscillator is called the CRY-dependent oscillator (CDO) system since it requires the blue-light photoreceptor CRYPTOCHROME (CRY) (Nsa *et al*. 2015).

In this work, we investigated the influence of short light-dark cycling conditions on the circadian clock in Neurospora, and compared the effects of the different light-dark cycles on growth, microconidia production, conidiation rhythmicity and gene expression. The findings of this work would shed new light on the knowledge of circadian clocks under conditions with extreme T-cycles.

## Materials and Methods

### Media, growth conditions and transformation procedure

The *Neurospora crassa 301-5* (*ras-1*^*bd*^, *a*) strain was used as the wild-type strain in this work. The 2% LCM media contained 1Vogel’s medium and 0.17% arginine, with 2% glucose. The race tube solid media consisted of 1Vogel’s medium containing 2% glucose, 50 ng of biotin/ml, and 1.5% agar.

The 315-13 (*fwd-1*^*RIP*^, his-3) strain, a strain in which the *fwd-1* was disrupted, was obtained from Dr Yi Liu’s lab and described previously (He *et al*. 2003). The *vvd*^*KO*^ strain was obtained from FGSC. All of the strains information are listed in supplemental Table S1.

Conidia were inoculated to seed mycelial mats in petri dishes, and disks were cut from these and used for 50-ml liquid cultures under certain LD conditions. At designated time points the cultures were harvested by filtration, frozen in liquid nitrogen, and ground in liquid nitrogen. Protein extraction, quantification and western blot analysis were conducted as described elsewhere (Görl *et al*. 2001). For western blotting, equal amounts of total protein (50 μg) were loaded in each protein lane of SDS-PAGE (7.5%) gels containing a ratio of 37.5:1 acrylamide/bisacrylamide.

An incubator (Percival Scientific, USA) was used to simulate different LD cycles, the white light intensity during light was 5000 lux or 1000 lux as indicated. The red light (λ= 660nm) was generated by an LED lamp, and the intensity was 5000 lux or 1000 lux as indicated. The blue light (λ= 470 nm) was generated by an LED lamp, and the intensity was 5000 lux or 1000 lux as indicated.

The counting of microconidia was conducted on a hemocytometer under an optical microscope. Basically, since the shape of microconidia is nearly round and they usually contain a single nucleus, we consider those hyphae with length/width ratio <1.5 as a subjective criteria as microconidia.

### Race tube assay

The conidiation banding profiles were assayed on race tubes under standard conditions (Nsa *et al*. 2015). The growth front was marked every 24 h under a red safe light in all light conditions in this work. All race tube experiments were carried out at room temperature (25°C).

### Protein analysis

Protein extraction, quantification, western blot analysis was performed as previously described (Guo *et al*. 2010).

### Dynamic modeling

See supplemental materials for details.

### RNA-seq and analysis

The strains subjected to RNA-seq were grown and the RNA samples were isolated from duplicates. All of the Pearson’s Correlation values between each duplicates were >0.99. Tophat was used as the aligner to map the reads to the reference genome [N. crassa OR74A (NC12)]. The genes showing ≥2-fold increase were selected for further analysis. RNA-seq data sets are available at the GEO database (GSE108814). See supplemental materials for more details.

### Statistical analysis

Data are mean ± SE or mean ± SD where indicated. n≥3. Student’s *t* test was used for all statistical analyses. * represents the p-value of the statistical tests is less than the significance level of 0.05 (p≤0.05); ** represents p≤ 0.01 and *** represents p≤ 0.001.

### Data availability

All strains and reagents are available upon request. The RNA-seq data sets are available at the GEO database (GSE108814). File S1 contains Table S1, Table S1, Figures S1-S5 and supplemental methods and protocols for dynamic modeling and RNA sequencing. Table S1 in File S1 lists all Neurospora strains. Table S2 in File S1 lists the parameters of the mathematical model. File S2 contains Table S3 that provides the up-regulated genes in *Δcry*, *Δvvd*, *Δwc-1* in 5000 lux. File S3 contains Table S4 that lists the KEGG pathway of genes differentially expressed in *Δcry*, *Δvvd*, *Δwc-1* in 5000 lux light and 1000 lux light. Figure S1 in File S1 shows the conidiation rhythms of FGSC4200 strain. Figure S2 in File S1 shows the FRQ protein levels under different LD routines. Figure S3 in File S1 provides the scheme for the mathematical model of *Neurospora* circadian rhythm. Figure S4 in File S1 shows the impact of light sensors on the adaptation to short LD cycles. Figure S5 in File S1shows the conidiation rhythms of *Neurospora* strains in short LD cycles. Figure S6 in File S1 shows the growth phenotype and conidiation rhythms of indicated strains.

## Results

### The synchronization and resonance of Neurospora rhythms under LD routines

To address how the short LD cycles affect the rhythms, we conducted race tube assays with the wild-type strain (*301-5, bd)*, under the following series of short LD cycling conditions including LD6:6, LD4:4, LD3:3, LD2:2, LD1:1, LD45min:45min and LD30min:30min, which consisted of symmetric light and dark phases. Additionally, we tested Neurospora growth in LD65min:25min, an asymmetric condition, to mimic the light/dark condition on orbital flight. The results show that under LD6:6, LD4:4, LD3:3, LD2:2, the conidiation displayed overt rhythms with periods that coincided with the LD cycles. However, when the cycles were shorter, under even shorter cycles, the entrained conidiation rhythms were abolished (Figure 1A).

**Figure 1.**
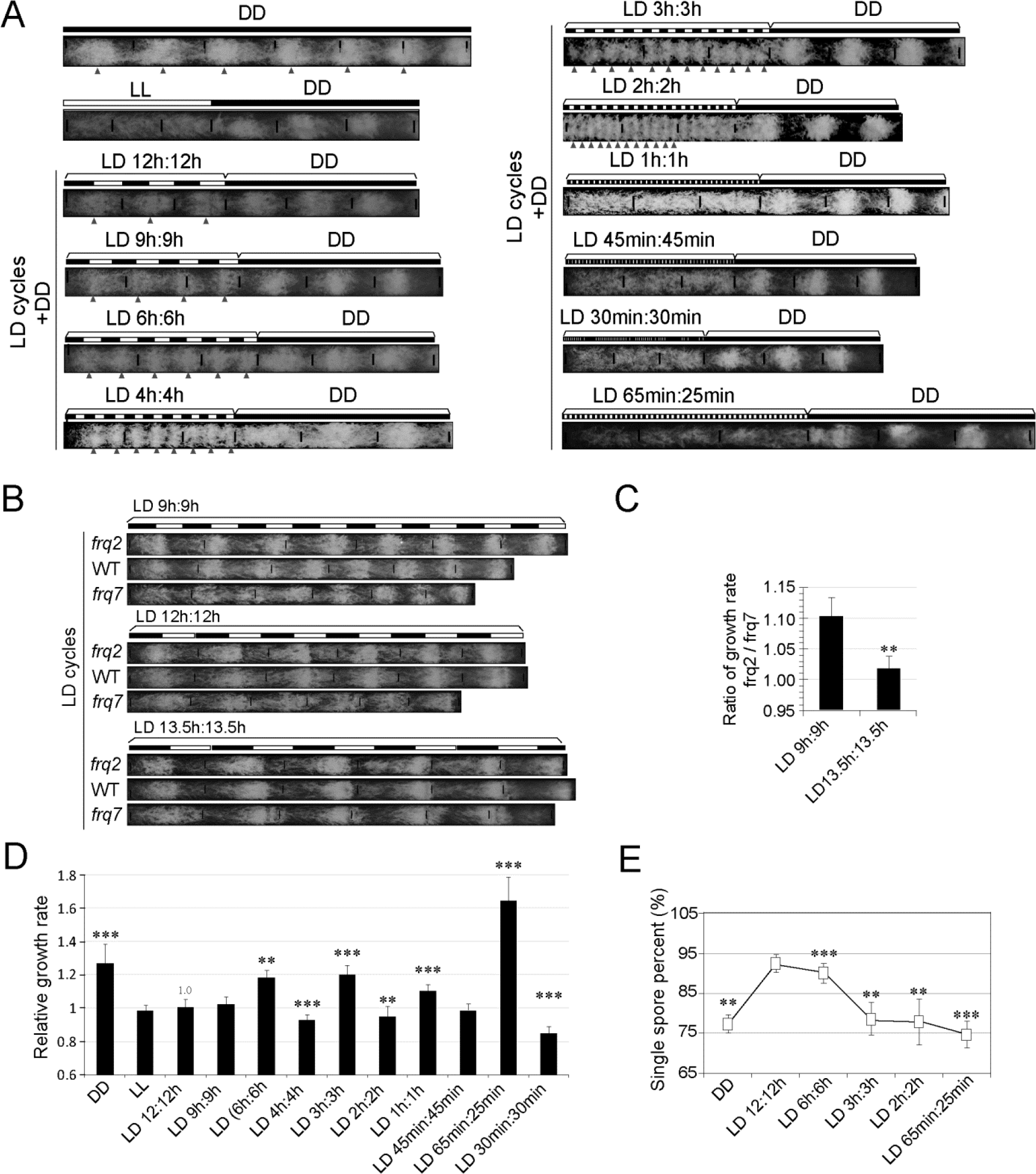
Conidiation rhythms, growth and adaptation of *Neurospora* in short LD cycles. A. Conidiation rhythms of *Neurospora* under a series of LD cycles. The black and white bars denote different LD regimes. Triangles denote the conidiation bands. B. Growth rate of *Neurospora* under a series of LD cycles. Data are mean ± SD, n=3. Represent results (n≥3) are shown. C. Ratio of the growth rates between *frq2* and *frq7*. Data are mean ± SD, n=3. D. Growth rates of indicated strains. The growth in LD12:12 was normalized to 1.0. Data are mean ± SE, n=3. E. Proportion of microconidia produced under different LD cycles. Data are mean ± SE, n=3.The light intensity was 5000 lux.

The arrhythmicity of Neurospora under extremely short LD cycles (e.g., LD1:1) could be explained by two possibilities: 1) the conidiation rhythms were abolished; 2) the bands were too compact and ambiguous to detect. To validate, we used the non-band strain FGSC4200 which grows very fast in race tube. In constant dark FGSC 4200 shows mild conidiation rhythms (Belden *et al*. 2007), while in LD1:1 it showed weak but recognizable conidiation rhythms (Figure S1 in File S1). These data suggest that although the free-running circadian period of Neurospora is approximately 22 h, the conidiation rhythm can be induced by extremely short LD cycles.

The circadian clock can adjust to fit with the cycling environment, especially the light. We compared the growth rate of two *frq* mutants *frq*^*2*^ and *frq*^*7*^, which possess shorter and longer free-running periods in constant dark, respectively (Gardner and Feldman 1981). The race tube assay results reveal that in LD9:9 the growth rate of *frq*^*2*^ was faster while in LD13.5:13.5 *frq*^*7*^ was faster (Figure 1, B and C), suggesting a resonance of Neurospora circadian period to the environment.

However, when we compared the growth rate of the wild-type strain under various LD cycles, we got unexpected results showing that Neurospora grew much faster in constant dark than that in constant light or most LD conditions. Moreover, the growth rate in shorter LD cycles was not necessarily slower than LD12:12. For instance, the growth in DD, LD6:6, LD3:3 and LD1:1 was significantly faster than that in LD12:12. The growth under LD65min:25min was also significantly faster than LD12:12 (Figure 1D).

Neurospora microconidia are small uninucleate spores that serve as male gametesor as asexual reproductive structures (Maheshwari 1999). When we counted the ratio of microconidia under LD12:12 and short LD cycles, respectively, and the results show that the ratio of microconidia in LD12:12 was significantly higher than those in other LD conditions (Figure 1E).

### Expression of the FRQ protein in different LD cycles

As the circadian clock is involved in controlling the conidiation rhythmicity, we asked whether the molecular rhythmicity of the FRQ protein is subject to changes in response to the length of the LD cycles. As one of the core factors in Neurospora circadian clock, FRQ acts as the negative component in the transcription/translational feedback loop (Baker *et al*., 2012). The western blotting results show that under LD12:12 and LD6:6, the expression of the FRQ protein and the phosphorylation profile displayed coincident rhythms with the periods of LD cycles. However, under LD3:3 and shorter cycles, the FRQ levels seemed to free run but we also observed some saw-tooth shaped fluctuations under LD3:3 (Figure 2, A-F and Figure S2 in File S1).

**Figure 2.**
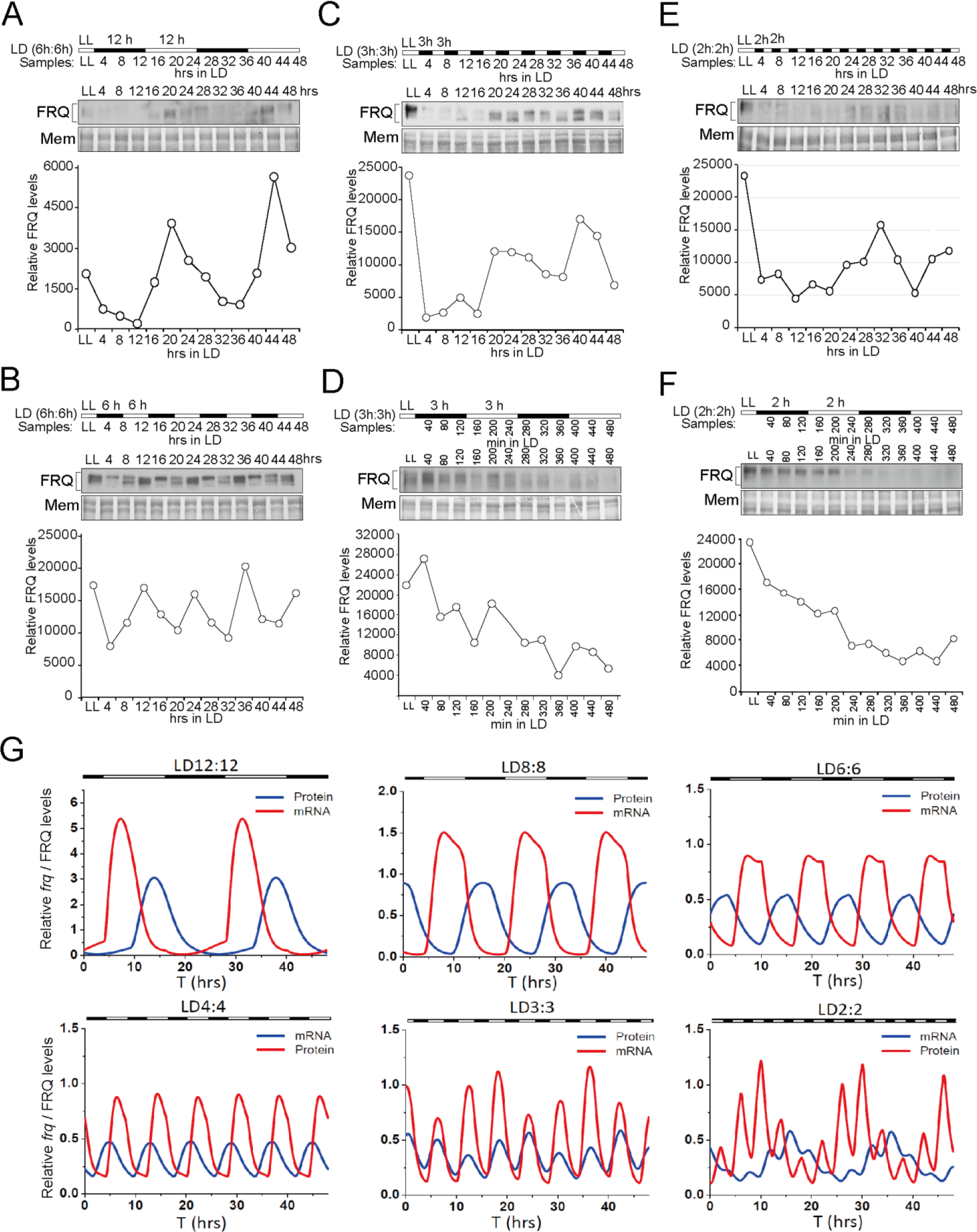
The FRQ protein levels under different LD routines. The FRQ protein levels are shown by western blot analysis. The LD routines are LD12:12 for two cycles (A), LD6:6 for 48 h (B), LD3:3 for 48 h (C), LD3:3for two cycles (D), LD2:2 for 48 h (E) and LD2:2 for two cycles (F), respectively. The black and white bars denote different LD regimes. Membranes (Mem) stained with Amido Black were used as loading control. The densitometric analysis of FRQ levels is shown at the bottom of each panel. Representative data of triplicates are shown. See more data in Fig. S1. The light intensity was 5000 lux.(G) Dynamic modeling results showing the time series of *frq* mRNA and FRQ protein under different light-dark cycles. Both the *frq* mRNA and FRQ protein can be entrained to 12:12 LD cycles, 8:8 LD cycles, 6:6 LD cycles and 4:4 LD cycles. As the frequency of the light signal increases to 3:3 cycles, the *frq* mRNA and FRQ protein still exhibit the oscillation with a period of approximately 6h. If the frequency of the light signal is further increased to 2:2 cycles, although the *frq* mRNA can respond to the induction by the light, the FRQ levels show the free running pattern with some fluctuations.

Mathematical modeling suggested that under LD12:12 and LD6:6, the concentration of FRQ protein matches the LD cycles (Figure 2G and Figure S3 in File S1 and Table S2 in file S1; supplemental protocol and parameters for modeling in File S1), instead, but underLD2:2 the FRQ levels show a free running pattern with some saw-tooth shaped fluctuations. The simulation results show that the mRNA levels have a direct response to light, but the protein curves are smoother, especially for high-frequency light signals. Therefore, the post-translational regulatory and translocation processes may perform a buffering function or high-frequency filtering function. Furthermore, the numerical data also predicted that the direct response of *frq* mRNA to the light stimuli is lower under shorter LD cycles (Figure 2G), which may be due to the light adaptation module (VVD-WCC loop) and the core negative feedback loop.

### FWO is not required for adaptation of conidiation rhythms to short LD cycles in high-intensity light

In the Neurospora circadian oscillator, WC-1 and WC-2 are the two positive elements in which WC-1 function as a blue light sensor (He *et al*. 2002). We asked whether WCC proteins are responsible for conidiation rhythmicity under LD cycles.

We grew the *wc-1*^*RIP*^, *wc-2*^*KO*^ and *wcc*^*DKO*^ and *frq*^*10*^ strains in race tubes under different LD cycles and observed condition rhythms for all of these mutants exhibit conidiation rhythms under LD12:12, LD6:6, LD3:3andLD2:2, but not in DD and LD45min:45min (Figure 3). The strains were exposed to a light intensity of 5000 lux, during the light periods. These data suggest that under short LD conditions, the conidiation rhythms cycle to the environment even without a functional circadian clock. We have also shown that in WT, the FRQ protein oscillation is masked under LD cycles (6≤T<24), but free runs when T<6 (Figure 2 and Figure S2 in File S1). Taken together, these results suggest that the FWO system is not required for the conidiation banding rhythms induced by the cycling environment.

**Figure 3.**
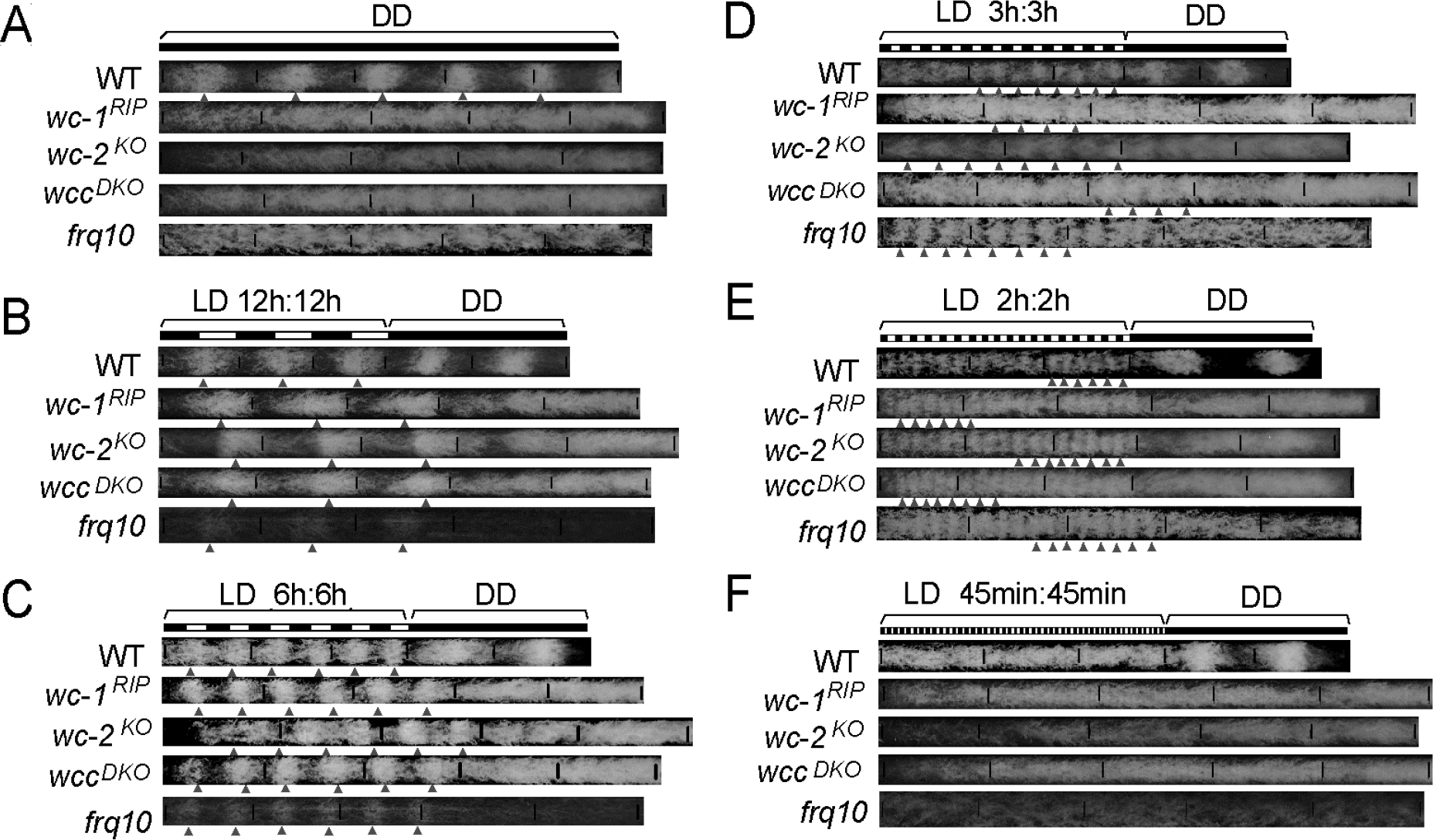
The conidiation rhythms of *Neurospora* strains in short LD cycles. The conditions were DD (A), LD12:12 (B), LD6:6 (C), LD3:3 (D), LD2:2 (E) and LD45min:45min (F). Represent results (n≥3) are shown. Triangles denote the conidiation bands. The strains were grown under white light and the light intensity was 5000 lux.

### Conidiation rhythmsof photosensor-related mutants under short LD regimes

First, we investigated the impacts of individual photosensing-associated genes on the conidiation rhythms under short LD cycles in Δ*cry*; Δ*vvd*; Δ*phy-1*; Δ*phy-2*; Δ*nop-1*and in *cog-1*, a newly identified mutant showing circadian conidiation rhythmicity under constant light (Nsa *et al*., 2015). WC-1, VVD and CRY are blue light sensors while PHY-1 and PHY-2 are potential red light receptors (Corrochano 2007; Froehlich *et al*. 2005; Chen *et al*. 2009). Unexpectedly, all of these strains showed conidiation rhythms with periods in accordance with the LD cycling periods, under conditions of LD12:12, LD6:6, LD3:3 and LD2:2 (Figure 4, A-D and Figure S4 in File S1). Together with the fact that the Δ*wc-1* strain can be entrained at LD2:2, these results suggest that either there exists additional photosensor(s), or the lack of an individual photosensor is not sufficient to abolish the adaptation of conidiation to short LD cycling cues.

**Figure 4.**
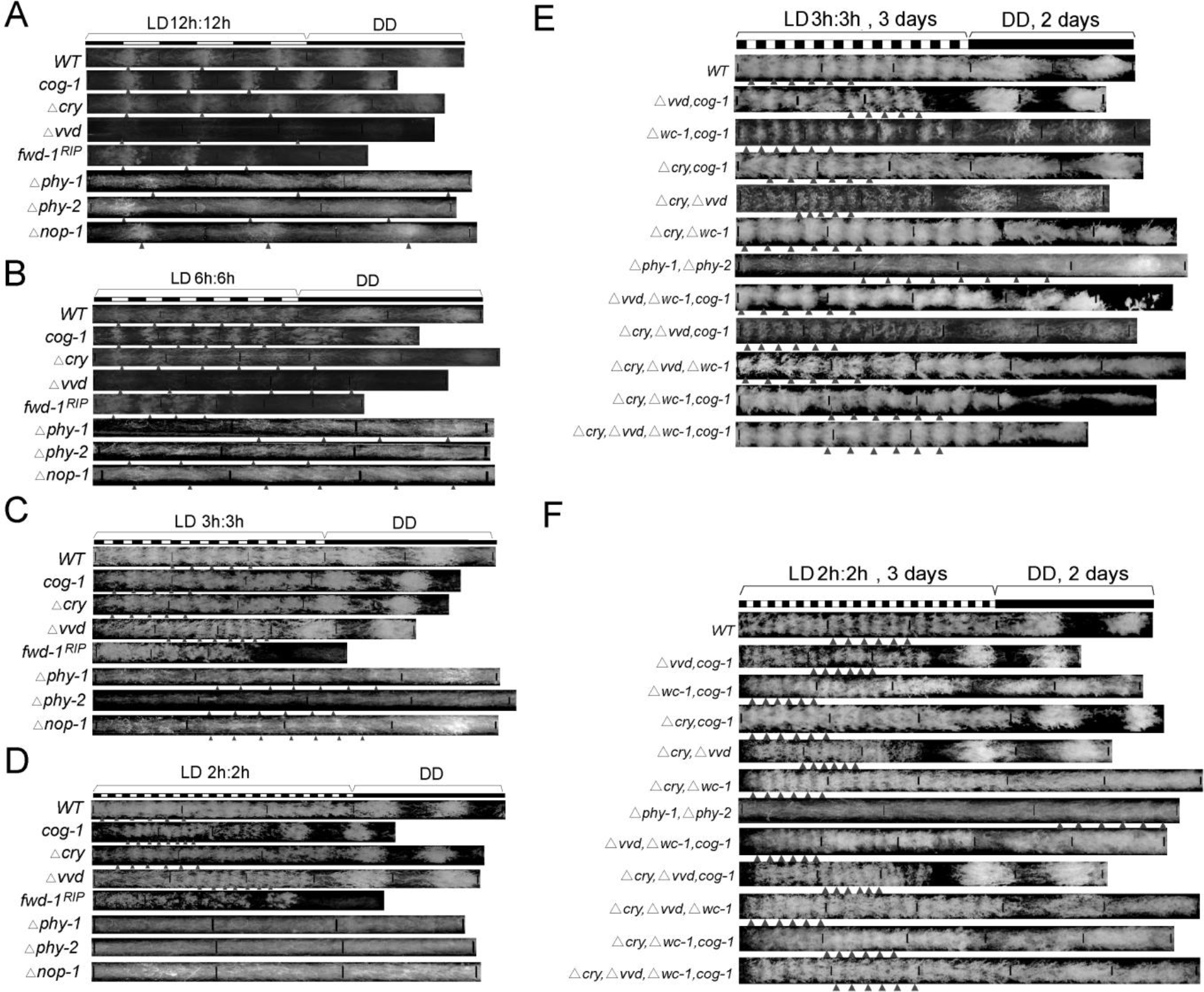
Impact of light sensors on the adaptation to short LD cycles. Race tube results of indicated strains under indicated LD cycles. Represent results (n≥3) are shown. Triangles denote the conidiation bands. The strains were grown under white light and the light intensity was 5000 lux.

With this in mind, we investigated the conidiation rhythms in strains bearing multiple mutations/deletions of light-sensor or related genes, which included Δ*cry*, Δ*vvd*; Δ*cry*, Δ*wc-1*; Δ*vvd*, *cog-1*; Δ*wc-1*, *cog-1*; Δ*vvd*, Δ*wc-1*, *cog-1*; Δ*cry*, Δ*vvd*, *cog-1*; Δ*cry*, Δ*vvd*, Δ*wc-1*; Δ*cry*, Δ*wc-1*, *cog-1* and Δ*cry*, Δ*vvd*, Δ*wc-1*, *cog-1*. Unexpectedly again, all of these strains still showed conidiation rhythms in short LD cycles (Figure 4, E and F).

We also observed these strains under~5000 lux red and blue light. In blue:dark (BD) cycles, most of the tested strains exhibit conidiation rhythms. In contrast, under red light, all the examined strains showed no entrained rhythms in LD6:6, LD3:3 and LD2:2 (Figure S5, A-G in File S1). Red light can also be sensed by some fungi, e.g., *Aspergillus nidulans*, *Trichoderma atroviride* and Neurospora, which is important for the asexual/sexual transition during development, hyphal growth and DNA stability (Corrochano 2007; Froehlich *et al*. 2005; Chen *et al*. 2009). Therefore, these data suggest that the conidiation rhythmicity is controlled predominantly by the blue light sensors and not the red light sensors.

The temperature increase that accompanied the switch-on of the lighting in the incubator, which might also have affected the conidiation. To rule out this possibility, we measured conidiation rhythms under temperature cycles (25.5°6h: 24.5°6h) as the range of incubator temperature is ~±0.5°C in our experiments, as previously described (Nsa *et al*. 2015). The results showed no synchronization in all tested strains (Figure S5H in File S1), demonstrating this temperature variation is not sufficient for synchronization. Taken together, these results show that all of these tested light sensors and associated factors were not critical for the conidiation rhythms. Instead, an unidentified blue light photoreceptor might be involved in controlling the conidiation rhythms.

The effects of different photosensing-associated genes on the production of carotenoid in all of these strains in constant light were also analyzed. As previously has shown, the strains containing *vvd* mutations, including Δ*vvd*; Δ*vvd*, *cog-1*; Δ*vvd*, Δ*cry* and Δ*vvd*, Δ*cry*, *cog-1* displayed bright orange color in their mycelia. In contrast, the mycelia color of strains Δ*vvd*; Δ*wc-1*, *cog-1*; Δ*cry*, Δ*vvd*, Δ*wc-1* and Δ*vvd*, Δ*cry*, Δ*wc-1*, *cog-1* were not bright orange. Interestingly, the *fwd-1*^*RIP*^ strain showed bright orange color similar to that of Δ*vvd* (Figure S6 in File S1).

In contrast to our results, Nsa *et al* showed that the Δ*wc-1* strain exhibit no obvious rhythms under a number of T-cycling conditions including LD6:6, LD9:9, LD12:12 and LD14:14 (Nsa *et al*. 2015). However, we used a white light intensity (~5000 lux) that was much higher than that used by Nsa et al (1200 lux), which may account for the inconsistency. In support of this view, when we grew the clock mutants in LD conditions and the light intensity was approximately 1000 lux, and conidiation rhythmicity was abolished at the tested LD conditions (Figure 5, A and B and Figure S5 in File S1 and Figure S6 in File S1). Conversely, the conidiation rhythms in clock gene mutants, *wc-1*^*RIP*^, *wcc*^*DKO*^, *frq*^*10*^ and Δ*wc-2*, were present in 5000 lux blue light but were absent in 1000 lux (Figure S6 in File S1). These data suggest that while clock components might act to promote the function of a potential photoreceptor to elicit conidiation rhythms, the response of clock mutants to a high-intensity but not low-intensity light, suggests that higher-intensity light might overcome the repression effect caused by the loss of clock components (Figure 5, C-E).

**Figure 5.**
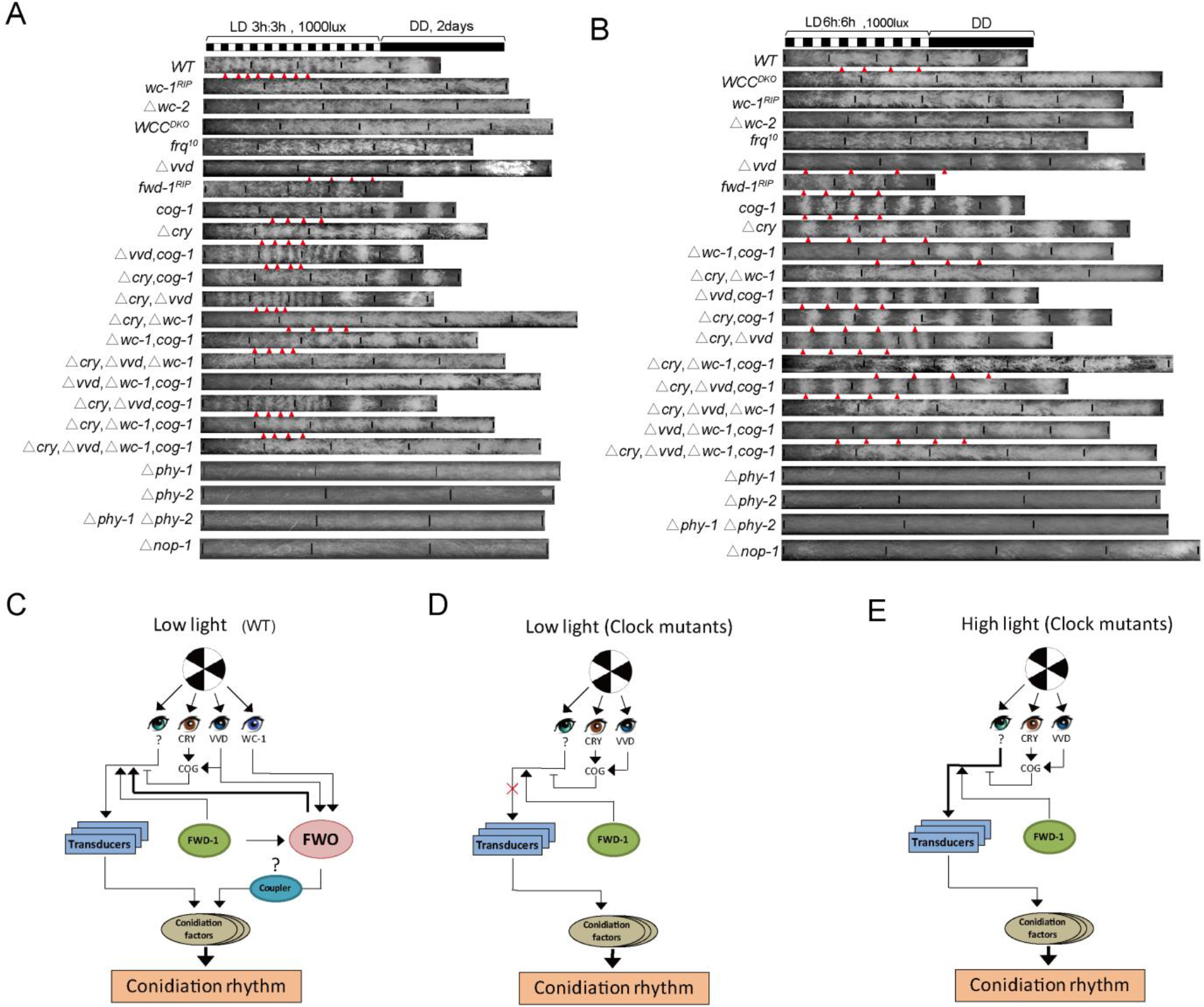
Conidiation rhythms in low white light (1000 lux). A/B. Race tube results of conidiation rhythms of indicated strains in LD3:3 (A) and LD6:6 (B). The white light intensity was 1000 lux. Triangles denote the conidiation bands. C-E. Schematics of the control of conidiation rhythms under short LD cycles of WT in low-intensity light (C) and clock mutants in low-intensity light (D) and high-intensity light (E). WC-1, VVD and additional photoreceptor(s)are implicated in the regulation of the conidiation rhythms. In 6 h≤ T ≤ 24 h, the FWO system is entrained, and the FRQ rhythms are not endogenous as they cannot be maintained in constant dark. The question masks denote unidentified photoreceptor and putative coupler linking FWO and conidiation, respectively. Transducers denote the factors linking photoreceptor and conidiation.

It is intriguing to note that in BD cycles (blue light: 1000 lux), the strains *cog-1*; Δ*wc-1*, *cog-1* and Δ*vvd*, Δ*wc-1*, *cog-1* exhibited biomodal patterns on the first day (Figure S5F in File S1), suggesting that the CDO pathway might play a role in mediating the synchronization of conidiation rhythmicity (Figure S5 in File S1).

### Implication of FWD-1 in regulating the conidiation rhythms under short LD cycles

To probe the possible mechanisms regulating the adaptation to LD cycles, we also checked the conidiation rhythms of several random mutants, among them the *fwd-1*^*RIP*^ strainthat lacks the functional *fwd-1* gene which encodes FWD-1, the Neurospora homolog of *Drosophila* Slimb, a ubiquitin E3 ligase. Neurospora FWD-1 has been implicated in the regulation of FRQ ubiquitination and turn over (He *et al*. 2003; Larrondo *et al*. 2015).

The *fwd-1*^*RIP*^ strain exhibited no overt conidiation bands under LD3:3 and shorter LD cycles (Figure 4 and Figure 5 and Figure S5 in File S1). The mutants lacking tested photosensors showed normal responses to short LD cycles, suggesting that the phenotype of *fwd-1*^*RIP*^ is not based on the impacts of FWD-1 on these genes, i.e., additional photosensor(s) controlled by FWD-1 might be implicated in regulating the conidiation rhythms under short LD cycles. Alternatively, a potential factor connecting the specific photosensor to conidiation and, that is controlled by FWD-1, might be implicated (Figure 5, C and D).

### Transcriptomic analysis of genes induced by high-intensity light

The above data suggest that high-intensity light influences conidiation rhythms, thus we conducted RNA-seq analysis to assess the changes of gene expression in response to high-intensity light.

We compared the transcriptomic changes of the strain Δ*cry*, Δ*vvd*, Δ*wc-1*, which contains no known blue light photoreceptors, after exposure for 45min to either 1000 lux or 5000 lux white light (Fig. 6A). The results showed that upon light exposure, a variety of genes involved in metabolism and gene expression were up-regulated (folds  1.5). For the genes exclusively induced by 5000-lux light, one putative motif (GAXGA)was identified by the XXmotif online tool (http://xxmotif.genzentrum.lmu.de/), which is present in 56 promoter areas of the 73 genes (Figure 6C). Light of 5000-lux induced the exclusive expression of 73 specific genes, many of which were enriched in fatty acid metabolism, ribosome biogenesis, etc. (Figure 6, C and D and Table S3 in File S2 and S4 in File S3). The induced expression of three representative genes (NCU00298, NCU00992 and NCU09771) were confirmed in Δ*cry*, Δ*vvd*, Δ*wc-1*; Δ*phy-1*, Δ*phy-2* and *Δnop-1* by qRT-PCR (Figure 6, E-G). Although the pathways of ribosome biogenesis and propanoate metabolism were induced in both 1000-lux and 5000-lux light (Table S4 in File S3), the genes involved were different, suggesting that higher-intensity light plays specific roles in regulating the expression certain genes and metabolism.

**Figure 6.**
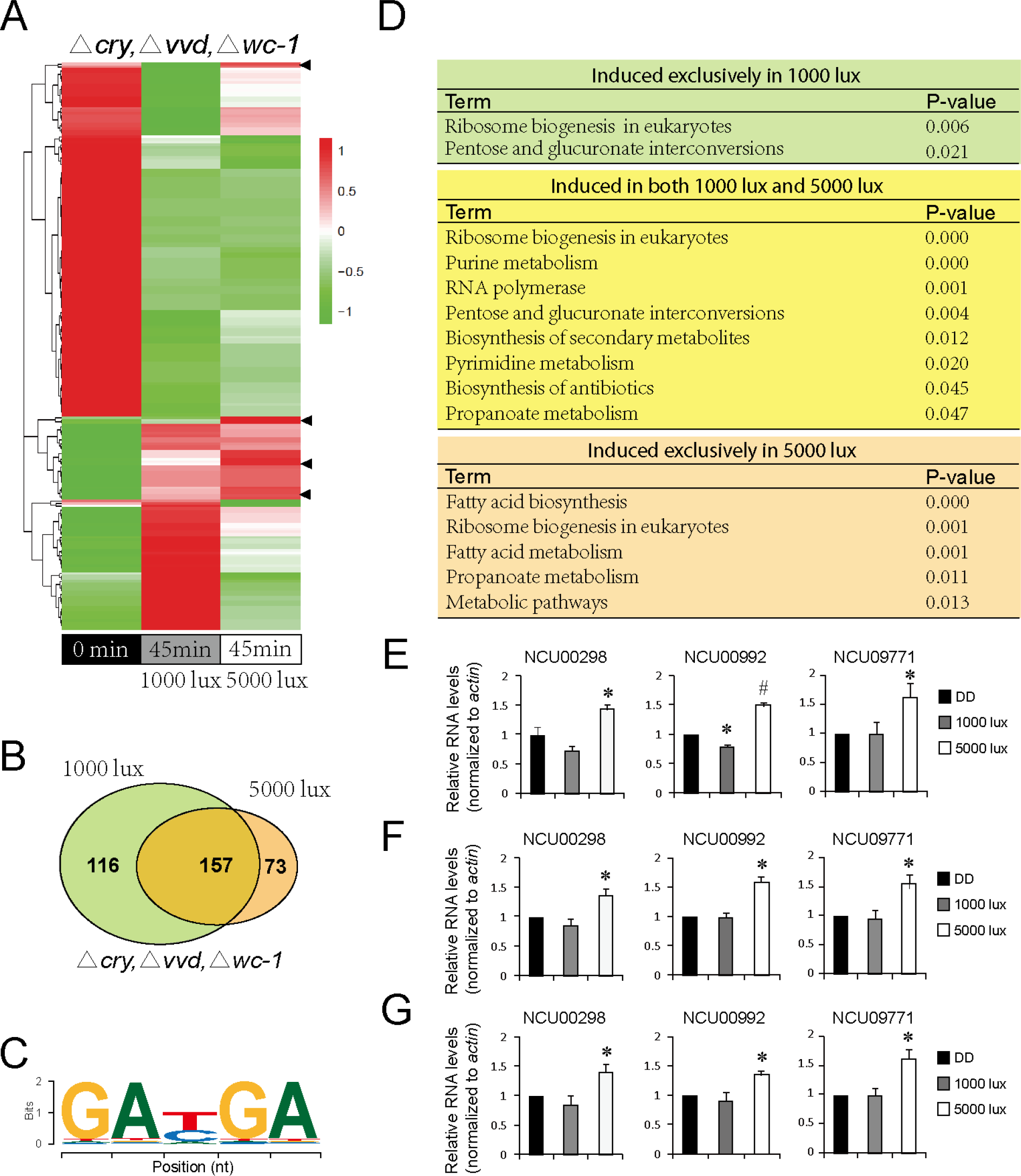
Characterization of genes showing light responses in high-intensity white light. A. Heat map of the differentially expressed genes in Δ*vvd,* Δ*cry,* Δ*wc-1* at DD24, 1000-lux white light and 5000-lux white light. The heat map was created according to the RNA-seq data. B. Venn diagram showing the genes up-regulated in 1000 lux and 5000 lux relative to DD24, respectively. C. Sequence logo of a predicted motif. D. Distribution of affected KEGG pathways according to the up-regulated genes revealed from RNA-seq. E-G. qRT-PCR validation of three genes specifically induced by 5000-lux light in Δ*vvd*, Δ*cry*, Δ*wc-1* (E), Δ*phy-1*, Δ*phy-2* (F) and Δ*nop-1* (G) strains. The gene expression was normalized to *actin*. The levels at DD24 was normalized to 1.0. Data are mean ± SE, n=3.

Taken together, these data confirm that high-intensity light induces the expression of a set of genes in clock mutants that might represent the targets for studying the potential unidentified photoreceptor(s).

## Discussion

Within a certain range, the cyclical environmental changes (*T*-cycles) can induce a rhythm with the same periodicity as the environment even when these are non-24-h cycles, which means *T*-cycles mask the endogenous circadian periodicity. This range varies in different organisms (Jud *et al*. 2005; Aschoff 1960). In Neurospora, we show that under a number of short LD conditions, including even very short cycles, such as LD1:1, induce conidiation banding rhythms with periods that change along with the environmental periods. Under constant darkness, Neurospora conidiation rhythms display an ~22 h period which reflects the endogenous circadian period. Our study shows that under almost all of the short LD cycles, the conidiation rhythms were induced by the *T*-cycles. The conidiation rhythms under short *T*-cycles are not endogenous as they disappeared in constant dark; instead, they are the hourglass-type rhythms.

FRQ is one of the central circadian clock components and the rhythmic changes in FRQ levels and phosphorylation patterns represent the endogenous circadian rhythms (Baker *et al*. 2012). When we further looked at the expression of the core circadian protein FRQ under short LD cycles, we found that similar to the conidiation rhythms, the period of FRQ changed in accordance to the LD cycles when *T*> LD3:3. In contrast, under LD cycles wit *T*≤ LD3:3, FRQ level no longer oscillated with the *T* cycles, and instead exhibit periods close to 24h, i.e., the masking effects faded while the FRQ rhythm free ran. The inconsistency between conidiation rhythms and FRQ rhythms might reflect the decoupling between the intrinsic and extrinsic rhythms under the extremely short LD cycles. The dynamic modeling results showed the superimposition of endogenous and the light-induced rhythms, which can also be observed in the western blot results of FRQ protein under very short LD cycles (*T* ≤LD3:3) (Figure 2 and Figure S2 in File S1).

In many species, it has been demonstrated that non-24h T-cycles decrease their fitness (Hut *et al*., 2011), suggesting the adaptation of the endogenous rhythm of an organism to external cycling cues is critical for the development and growth. We show that the growth rate of Neurospora was slower under LD12:12 compared to many other conditions. However, the proportion of microconidia produced in LD12:12 prevailed over those in other conditions.

Faster growth of an organism does not necessarily mean optimal fitness to the environment (Hut *et al*. 2011). Instead, the capability to produce variable and fertile progenies may be more crucial. Neurospora microconidia function predominantly as spermatid (Maheshwari 1999), suggesting that the LD12:12 condition might be optimal for sexual reproduction. It is likely that in nature, the circadian clock renders Neurospora better prepared to use its sexual reproduction strategies to cope with constantly changing environmental stresses. Compared to the findings in some other organisms, these results reflect that different organisms may employ different strategies for adaptation and propagation (Vaze and Sharma 2013). In addition, the dramatic differences in growth under LL vs DD, LD45min:45min vs LD65min:25min indicate that the growth is influenced by the daily length of lighting in Neurospora, which might be similar to the seasonal physiology responses in different kingdoms (Arendt 1988; Tan *et al*. 2004).

In fungi, a number of developmental and physiological processes are affected by light (Lauter 1996; Linden *et al*. 1997). In Neurospora, the most prominent light response is the light-regulated biosynthesis of the photo protective mycelia carotenoids (Schrott 1980; Harding and Turner 1981), phototropism of perithecial beaks, protoperithecia development, promotion of conidia, light entrainment of the circadian clock and the formation of spores and phototropism (Lauter *et al*. 1997; Ballario *et al*. 1996).Almost all Neurospora light responses described so far are only known to be triggered by blue light. Transcriptomic analysis showed that expression of some genes can be triggered by light in the *wc-1* null strain, suggesting the existence of additional photoreceptors (Chen *et al*. 2009). Surprisingly, we show that upon removal of several photosensors, the conidiation rhythms still exist in short LD cycles. In addition, the strains bearing *cog-1* mutation can still be entrained (Figure 3 and Figure 4 and Figure S4 and Figure S5 in File S1). Together with the results of cog-1 in BD3:3 (Figure S5F in File S1), these data suggest that the conidiation rhythms in short LD cycles are independent of either FWO or CDO although the CDO system affects the conidiation rhythms in short LD cycles.

In Neurospora, in addition to WC-1, there are several additional confirmed or putative photoreceptors in Neurospora including VVD, CRY, PHY1/2 and NOP-1 (Chen *et al*. 2009; Chen *et al*. 2010a; Chen and Loros 2009). VVD is a small LOV domain-containing blue-light photoreceptor protein that affects photo adaptation for many light-responsive genes (Zoltowski *et al*. 2007; Schwerdtfeger and Linden 2003). As has been known, some light sensors can regulate other light sensors at transcriptional or post-transcriptional levels. For instance, in the light, WCC regulates the expression of VVD in Neurospora. Conversely, VVD inhibits WCC in a light-dependent fashion though it is not essential for clock function. Furthermore, VVD has been shown to physically interact with WCC (Zoltowski *et al*. 2007; Chen *et al*. 2010a; Malzahn *et al*. 2010; Gin *et al*. 2013; Shrode *et al*. 2001). Cryptochrome is a putative blue light sensor and its function remains elusive (Froehlich *et al*. 2005; Chen *et al*. 2009). PHY1/2 are putative red light receptors and NOP-1 is a putative green light receptor (Chen *et al*. 2010b).

It has been claimed that WCC and VVD are responsible for most light-dependent physiological processes, if not all. However, the clock mutants showed synchronization to 5000 lux white light but not 1000 lux white light (Nsa *et al*. 2015) (Figure 3 and Figure 5). Moreover, the conidiation rhythmicity only occurs in blue light but not red light (Figure S5 in File S1).

Collectively, these data suggest the presence of an unidentified blue photoreceptor which induces conidiation rhythmicity. Interestingly, the synchronization of conidiation rhythms is independent of the FWO system; instead, the circadian clock might promote the conidiation responses (Figure 5, C and D).

Under 1000 lux white light, clock mutants displayed no conidiation rhythms, suggesting that the circadian clock functions to maintain or promote conidiation responses to short LD cycles. These data also suggest that the circadian clock might help Neurospora to promptly adjust its conidiation according to changeable light condition in the natural environment; for instance, caused by cloud movement or shadows being cast in the daytime. Under high-intensity light, even the clock mutants showed conidiation rhythms, suggesting the unidentified photoreceptor(s) might only sense very strong light, which can confer the condition rhythmicity in clock mutants.

Light initiates adaptation in a comprehensive set of metabolic pathways (Tisch and Schmoll 2010). In this work, we found that a subset of metabolic pathways was affected upon higher light exposure relative to lower light, suggesting that additional photoreceptor(s) than those that are known (e.g., WC-1, VVD and CRY) play a role in regulating light-induced metabolic adaptation.

FWD-1 is another regulator of conidiation rhythms in short LD cycles. In LD cycles (6h≤*T*<12h), the depletion of FWD-1 showed no overt impacts on conidiation rhythms, suggesting that FWD-1 does not affect the NFLO pathways. However, in shorter LD cycles (*T*<6h), the depletion of FWD-1 caused abolishment of conidiation rhythmicity, suggesting that FWD-1 controls NFLO pathways under such conditions. FWD-1 might function to affect the responsible photoreceptor(s) or sensitize transducers that couple specific light sensors and conidiation. Therefore, a lack of FWD-1 leads to abolishment of conidiation rhythms responding to the very short LD cycles.

The light conditions in certain extreme environments, e.g., space, dramatically differ from that on Earth. In low orbit, the LD cycles are very short (90-120-minute) with only 30% of the time in light and the remainder of the time in the dark (Stampi 1994). The present data also demonstrate that the circadian rhythms and fitness is impaired in simulated LD cycles with a 90-min period (Figure 1).

## Acknowledgements

We thank Prof. Deborah Bell-Pedersen (Texas A&M University, College Station, Texas, USA), Prof. Jay C. Dunlap (Geisel School of Medicine at Dartmouth). We thank Prof. Deborah Bell-Pedersen and Prof. Shaojie Li (Institute of Microbiology, CAS, China) for inspiring discussion and suggestions. This work was supported by the National 973 Program of China (Nos.2011CB711000 and 2012CB947600), National Natural Science Foundation of China (Nos. 31571205 and 31171119).

## Literature cited

Abraham, U., A. E. Granada, P. O. Westermark, M. Heine, A. Kramer et al., 2010 Coupling governs entrainment range of circadian clocks. Mol. Syst. Biol. 6: 438.

Arendt, J., 1998 Melatonin and the pineal gland: influence on mammalian seasonal and circadian physiology. Rev. Reprod. 3: 13–22.

Aronson, B.D., K. A. Johnson, J. J. Loros, and J. C. Dunlap, 1994 Negative feedback defining a circadian clock: autoregulation of the clock gene frequency. Science 263: 1578–1584.

Aschoff, J., 1960 Exogenous and endogenous components in circadian rhythms. Cold Spring Harb. Symp. Quant. Biol. 25: 11–28.

Baker, C.L., J. J. Loros, and J. C. Dunlap, 2012 The circadian clock of Neurospora crassa. Fems. Microbiol. Rev. 36: 95–110.

Ballario, P., P. Vittorioso, A. Magrelli, C. Talora, and A. Cabibbo et al., 1996 White collar-1, a central regulator of blue light responses in Neurospora, is a zinc finger protein. EMBO J. 15: 1650–1657.

Belden, W.J., L. F. Larrondo, A. C. Froehlich, M. Shi, C. Chen et al., 2007 The band mutation in Neurospora crassa is a dominant allele of ras-1 implicating RAS signaling in circadian output. Genes Dev. 21: 1494–1505.

Bell-Pedersen, D., V. M. Cassone, D. J. Earnest, S. S. Golden, P. E. Hardin et al., 2005 Circadian rhythms from multiple oscillators: Lessons from diverse organisms. Nat. Rev. Genet. 6: 544–556.

Binkley, S. A., 1990 The clockwork sparrow: time, clocks, and calendars in biological organisms. Prentice-Hall, Inc., New Jersey.

Boulos, Z., M. M. Macchi, and M. Terman, 2002 Twilights widen the range of photic entrainment in hamsters. J. Biol. Rhythms 17: 353–363.

Chen, C., and J. J. Loros, 2009 Neurospora sees the light: light signaling components in a model system. Commun Integr Biol 2:448–451.

Chen, C., B. S. Demay, A. S. Gladfelter, J. C. Dunlap, and J. J. Loros, 2010 Physical interaction between VIVID and white collar complex regulates photoadaptation in Neurospora. Proc. Natl. Acad. Sci. USA 107: 16715–16720. a

Chen, C., J. C. Dunlap, and J. J. Loros, 2010 Neurospora illuminates fungal photoreception. Fungal Genet. Biol. 47: 922–929. b

Chen, C., C. S. Ringelberg, R. H. Gross, J. C. Dunlap, and J. J. Loros, 2009 Genome-wide analysis of light-inducible responses reveals hierarchical light signalling in Neurospora. EMBO J. 28: 1029–1042.

Corrochano, L. M., 2007 Fungal photoreceptors: sensory molecules for fungal development and behaviour. Photochem. Photobiol. Sci. 6: 725–736.

Czeisler, C.A., J. F. Duffy, T. L. Shanahan, E. N. Brown, J. F. Mitchell et al., 1999 Stability, precision, and near-24-hour period of the human circadian pacemaker. Science 284: 2177–2181.

Dong, W., X. Tang, Y. Yu, R. Nilsen, R. Kim et al., 2008 Systems biology of the clock in Neurospora crassa. Plos One 3: e3105.

Dunlap, J. C., J. J. Loros, J.J., and P. J. DeCoursey, 2004 Chronobiology: biological timekeeping. Sinauer Associates. Inc. Publishers.

Froehlich, A. C., Y. Liu, J. J. Loros, and J. C. Dunlap, 2002 White Collar-1, a circadian blue light photoreceptor, binding to the frequency promoter. Science 297: 815–819.

Froehlich, A.C., B. Noh, R. D. Vierstra, J. J. Loros, and J. C. Dunlap, 2005 Genetic and Molecular Analysis of Phytochromes from the Filamentous Fungus Neurospora crassa. Eukaryot. Cell 4: 2140–2152.

Gardner, G. F., and J. F. Feldman, 1981 Temperature compensation of circadian period length in clock mutants of Neurospora crassa. Plant Physiol. 68: 1244–1248.

Gin, E., A. C. Diernfellner, M. Brunner, and T. Höfer, 2013 The Neurospora photoreceptor VIVID exerts negative and positive control on light sensing to achieve adaptation. Mol. Syst. Biol. 9: 667.

Görl, M., M. Merrow, B. Huttner, J. Johnson, T. Roenneberg et al., 2001 A PEST‐like element in FREQUENCY determines the length of the circadian period in Neurospora crassa. EMBO J. 20: 7074–7084.

Guo, J., G. Huang, J. Cha, and Y. Liu, 2010 Biochemical methods used to study the gene expression and protein complexes in the filamentous fungus Neurospora crassa. Methods Mol. Biol. 638: 189–200.

Harding, R. W., and R. V. Turner, 1981 Photoregulation of the carotenoid biosynthetic pathway in albino and white collar mutants of Neurospora crassa. Plant Physiol. 68: 745–749.

He, Q., P. Cheng, Y. Yang, Q. He, H. Yu et al., 2003 FWD1-mediated degradation of FREQUENCY in Neurospora establishes a conserved mechanism for circadian clock regulation. EMBO J. 22: 4421–4430.

He, Q., P. Cheng, Y. Yang, L. Wang, and K. H. Gardner et al., 2002 White collar-1, a DNA binding transcription factor and a light sensor. Science 297: 840–843.

Heintzen, C., J. J. Loros, and J. C. Dunlap, 2001 The PAS protein VIVID defines a clock-associated feedback loop that represses light input, modulates gating, and regulates clock resetting. Cell 104: 453–464.

Highkin, H.R., and J. B. Hanson, 1954 Possible interaction between light-dark cycles and endogenous daily rhythms on the growth of tomato plants. Plant Physiol. 29: 301–302.

Hurley, J. M., J. J. Loros, and J. C. Dunlap, 2015 Dissecting the mechanisms of the clock in Neurospora. Methods Enzymol. 551: 29–52.

Hut, R. A., and D. G. Beersma, 2011 Evolution of time-keeping mechanisms: early emergence and adaptation to photoperiod. Philos. Trans. R. Soc. Lond. B Biol. Sci. 366: 2141–2154.

Jewett, M. E., R. E. Kronauer, and C. A. Czeisler, 1994 Phase-amplitude resetting of the human circadian pacemaker via bright light: a further analysis. J.Biol. Rhythms 9: 295–314.

Jud, C., I. Schmutz, G. Hampp, H. Oster, and U. Albrecht, 2005 A guideline for analyzing circadian wheel-running behavior in rodents under different lighting conditions. Biol. Proced. Online 7: 101–116.

Larrondo, L.F., C. Olivaresyanez, C. L. Baker, J. J. Loros, and J. C. Dunlap, 2015 Decoupling circadian clock protein turnover from circadian period determination. Science 347: 1257277.

Lauter, F., 1996 Molecular genetics of fungal photobiology. J. Genet. 75: 375–386.

Lauter, F., C. T. Yamashiro, and C. Yanofsky, 1997 Light stimulation of conidiation in Neurospora crassa: studies with the wild-type strain and mutants wc-1, wc-2 and acon-2. J. Photochem. Photobiol. B 37: 203–211.

Lewis, P. R., and M. C. Lobban, 1957 Dissociation of diurnal rhythms in human subjects living on abnormal time routines. Exp. Physiol. 42: 371–386.

Linden, H., M. Rodriguez-Franco, and G. Macino, 1997 Mutants of Neurospora crassa defective in regulation of blue light perception. Mol. Gen. Genet. 254: 111–118.

Madrid, J., F. Sanchez-Vazquez, P. Lax, P. Matas, E. Cuenca et al., 1998 Feeding behavior and entrainment limits in the circadian system of the rat. Am. J. Physiol. 275: R372–R383.

Maheshwari, R., 1999 Microconidia of Neurospora crassa. Fungal Genet. Biol. 26: 1–18.

Malzahn, E., S. Ciprianidis, K. Káldi, T. Schafmeier, and M. Brunner, 2010 Photoadaptation in Neurospora by competitive interaction of activating and inhibitory LOV domains. Cell 142: 762–772.

Nsa, I. Y., N. Karunarathna, X. Liu, H. Huang, B. Boetteger, et al., 2015 A novel cryptochrome-dependent oscillator in Neurospora crassa. Genetics 199: 233–245.

Ouyang, Y., C. R. Andersson, T. Kondo, S. S. Golden, and C. H. Johnson, 1998 Resonating circadian clocks enhance fitness in cyanobacteria. Proc. Natl. Acad. Sci. USA 95: 8660–8664.

Ralph, M. R., and M. Menaker, 1988 A mutation of the circadian system in golden hamsters. Science 241: 1225–1227.

Refinetti R., 2004 Parameters of photic resetting of the circadian system of a diurnal rodent, the Nile grass rat. Acta. Sci. Vet. 32: 1–6.

Schwerdtfeger, C., and H. Linden, 2001 Blue light adaptation and desensitization of light signal transduction in Neurospora crassa. Mol. Microbiol. 39: 1080–1087.

Shrode, L. B., Z. A. Lewis, L. D. White, D. Bellpedersen, and D. J. Ebbole, 2001 vvd is required for light adaptation of conidiation-specific genes of Neurospora crassa, but not circadian conidiation. Fungal Genet. Biol. 32: 169–181.

Schrott, E. L., 1980 Fluence response relationship of carotenogenesis in Neurospora crassa. Planta 150: 174–179.

Schwerdtfeger, C., and H. Linden, 2003 VIVID is a flavoprotein and serves as a fungal blue light photoreceptor for photoadaptation. EMBO J. 22: 4846–4855.

Stampi, C., 1994 Sleep and circadian rhythms in space. J.Clin. Pharmacol. 34: 518–534.

Tan, Y., M. Merrow, and T. Roenneberg, 2004 Photoperiodism in Neurospora crassa. J. Biol. Rhythms 19: 135–143.

Tisch, D., and M. Schmoll, 2010 Light regulation of metabolic pathways in fungi. Appl. Microbiol. Biotechnol. 85:1259–1277.

Vaze, K.M., and V. K. Sharma, 2013 On the adaptive significance of circadian clocks for their owners. Chronobiol. Int. 30: 413–433.

Zoltowski, B.D., C. Schwerdtfeger, J. Widom, J. J. Loros, A. M. Bilwes et al., 2007 Conformational switching in the fungal light sensor Vivid. Science 316: 1054–1057.

